# A multi-layer mean-field model for the cerebellar cortex: design, validation, and prediction

**DOI:** 10.1101/2022.11.24.517708

**Authors:** Roberta M. Lorenzi, Alice Geminiani, Yann Zerlaut, Alain Destexhe, Claudia A.M. Gandini Wheeler-Kingshott, Fulvia Palesi, Claudia Casellato, Egidio D’Angelo

## Abstract

Mean-field (MF) models can be used to summarize in a few statistical parameters the salient properties of an inter-wired neuronal network incorporating different types of neurons and synapses along with their topological organization. MF are crucial to efficiently implement the modules of large-scale brain models maintaining the specificity of local microcircuits. While MFs have been generated for the isocortex, they are still missing for other parts of the brain. Here we have designed and simulated a multi-layer MF of the cerebellar network (including Granule Cells, Golgi Cells, Molecular Layer Interneurons, and Purkinje Cells) and validated it against experimental data and the corresponding spiking neural network (SNN) microcircuit model. The cerebellar MF was built using a system of equations, where properties of neuronal populations and topological parameters are embedded in inter-dependent transfer functions. The model time constant was optimised using local field potentials recorded experimentally from acute mouse cerebellar slices as a template. The MF satisfactorily reproduced the average dynamics of the different neuronal populations in response to various input patterns and predicted the modulation of Purkinje Cells firing depending on cortical plasticity, which drives learning in associative tasks, and the level of feedforward inhibition. The cerebellar MF provides a computationally efficient tool that will allow to investigate the causal relationship between microscopic neuronal properties and ensemble brain activity in virtual brain models addressing both physiological and pathological conditions.

## 1. Introduction

Brain modelling is opening new frontiers for experimental and clinical research toward personalised and precision medicine (Amunts et al., 2013; Schirner et al., 2015). Brain models can be developed at different scale, ranging from microscopic properties of neurons and microcircuits to the ensemble behaviour of the whole brain. Arguably, a model spanning across scales would increase the fidelity in modelling single brain regions, improving the accuracy of whole-brain dynamics simulations (D’Angelo and Jirsa, 2022), but this clearly bears conceptual and practical drawbacks. At the microscale, Spiking Neural Networks (SNNs) reproduce neural circuits as a set of interconnected neurons (Plesser et al., 2015; Yavuz et al., 2016; Knight et al., 2021): the state of each neuron and synapse in the network is updated at each simulation step, allowing to investigate neural circuits functioning at a high level of granularity and biological plausibility. However, this degree of detail is hard to manage when simulating brain signals, like those derived from electroencephalography (EEG) or functional magnetic resonance imaging (fMRI). To manage the high complexity of brain signals, the dynamics of a neuronal population have been condensed into ensemble density models called neural masses. These provide a description of the expected values of neuronal activity states, under the assumption that the equilibrium density has a point mass (Wilson and Cowan, 1972; Jansen and Rit, 1995). Neural fields are obtained from neural mass models when considering spatial information: these can be used to model spatial propagation of activity throughout brain volumes (Deco et al., 2008). Despite being computationally efficient and easy to fit on brain signals data, neural mass and neural field models lack a direct link to the microscopic scale, a fact that limits their applicability in investigating the neuronal bases of brain dynamics and the causal relationships between neural mechanisms at different scales.

A different approach is based on the mean-field (MF) approximation. The MF theory provides a general formalism to approximate high-dimensional random models by averaging the original system properties over degrees of freedom, i.e., maintaining the first two statistical moments (mean and variance) of the system. In neuroscience, MFs have been used to provide a representation of neuronal population dynamics, by replacing multiple single-neuron input-output (I/O) relationship with one based on the MF of the interconnected populations. MFs thus summarize the neuronal and connectivity properties of an entire spiking microcircuit through *ad-hoc* transfer functions (TFs) (Amit and Brunel, 1997; Brunel and Sergi, 1998; Kumar et al., 2008) and capture the statistical properties of network activity by computing the probabilistic evolution of neuronal states at subsequent time intervals (Kuhn et al., 2004; El Boustani and Destexhe, 2009; Zerlaut et al., 2016).

MFs can be used to investigate macroscale phenomena, such as brain rhythms and coherent oscillations (Cakan and Obermayer, 2020), and are computationally advantageous, with increased computational speed and low memory requirements compared to SNNs. Among current limitations, MFs do not capture in full the complex properties of specific neuronal populations and are valid only in certain firing regimes, e.g., at low frequency (Carlu et al., 2020).

Moreover, while a diversification of MFs for specific cortical regions has been proposed (Marreiros et al., 2008; Bastos et al., 2012; Deco et al., 2014; Auksztulewicz and Friston, 2015; Glomb et al., 2017; El Houssaini et al., 2020; Naskar et al., 2021), few attempts to develop MFs for subcortical regions have been performed (Moran et al., 2011; Saggar et al., 2015; van Wijk et al., 2018; Levenstein et al., 2019), despite their fundamental role in controlling brain dynamics and behavior (Schutter and van Honk, 2006; Castellazzi et al., 2014, 2018; Casiraghi et al., 2019; Andersen et al., 2020). In particular, the cerebellum has a dense connectivity with the cerebral cortex and remarkably impacts on whole-brain dynamics in resting-state and task-dependent fMRI (Casiraghi et al., 2019; Palesi et al., 2020; Monteverdi et al., 2022) prompting for the development of specific MFs to be included into whole-brain simulators.

The cerebellar cortex receives inputs from mossy fibers and climbing fibers and sends outputs to the deep cerebellar nuclei. Granule Cells (GrC), Golgi Cells (GoC), Molecular Layer Interneuron (MLI) and Purkinje Cells (PC) constitute the backbone of the cerebellar cortex, which shows a peculiar anisotropic geometry implementing a forward architecture with limited lateral connectivity and recurrent excitation. These properties, along with the neuronal types, differ remarkably from those of the cerebral cortex prompting for the definition of a specific MF. This operation represents both a challenge and an opportunity. All the neuron types of the cerebellar cortex, following a careful characterization in electrophysiology experiments in rodents *in vitro* and *in vivo* (D’Angelo et al., 2001; McKay and Turner, 2005; Molineux et al., 2006; Solinas et al., 2007; Lachamp et al., 2009), have been represented by detailed multicompartmental models (Masoli et al., 2015, 2020b, 2020a; Masoli and D’Angelo, 2017; Rizza et al., 2021), simplified into point-neuron models (Lennon et al., 2014; Geminiani et al., 2018, 2019a), and embedded in network models of the cerebellar microcircuit (Solinas et al., 2007; Geminiani et al., 2019b, 2019a; Casali et al., 2020; De Schepper et al., 2022). Thus, the cerebellum provides an ideal substrate for generating a MF, in which the internal dynamics can be remapped onto a precise physiological counterpart and validated against a rich and informative dataset.

In this work we have developed and validated a multi-layer MF of the cerebellar cortex, which maintains the salient properties of the inter-wired cerebellar neuronal populations. Indeed, the MF was derived from a biology-grounded model of the cerebellar microcircuit, which was used to define the topology and tune the parameters of the MF, and it was then validated against a rich set of SNN outputs. The perspective is to integrate the present mesoscopic cerebellar MF as a module of macroscale models, e.g. Dynamic Causal Modeling (DCM) (Friston et al., 2003, 2019; Parr et al., 2020) or The Virtual Brain (TVB)(Sanz Leon et al., 2013), to simulate the cerebellar contribution to brain activity in physiological and pathological conditions.

## 2. Methods

In this section we describe the development, tuning, validation, and application of a multi-layer MF of the cerebellar circuit (Figure 1). The MF formalism provides a statistical summary of a SNN activity through the first two statistical moments (i.e. average and variance) of the population firing rates (El Boustani and Destexhe, 2009). Here the SNN bottom-up modelling approach is merged with the standard MF mathematical formalism to obtain a multi-layer MF of the cerebellar cortex. In order, we present the cerebellar SNN model used as the structural and functional reference of the MF (**section 2.1**), the design of the MF architecture based on cerebellar topology (**section 2.2.1**), the implementation of the MF equations derived with an heuristic approach (**section 2.2.2**) (Zerlaut et al., 2016, 2018; Carlu et al., 2020), the protocols used to optimise the MF time constant (**section 2.2.3**) and to validate the MF (**section 2.3**), the applications of the MF to predict the activity modulation induced by different levels of synaptic plasticity and of inhibitory control (**section 2.2.4**).

**Figure 1.**
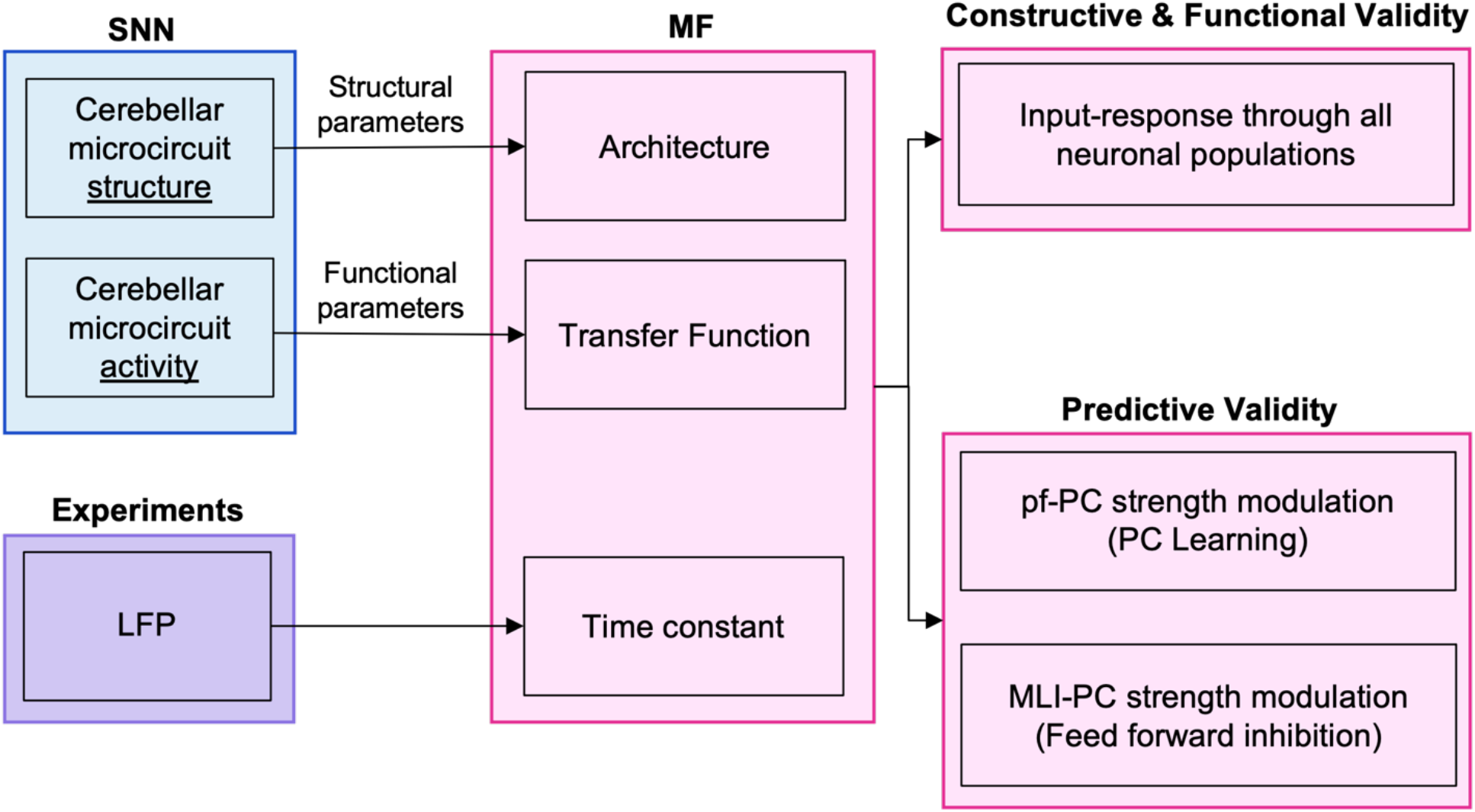
Pipeline of the multi-layer cerebellar MF model. The workflow of the study is represented. MF was designed based on structural and functional parameters extracted from Spiking Neural Network (SNN) simulations. The time constant of the resulting MF was optimized against Local Field Potential (LFP) experimental data. The model was first validated against neural activity of SNN with different stimulation protocols and then used to reproduce the effect of synaptic plasticity in molecular layer interneurons.

### 2.1 SNN model

This cerebellar cortex model was built using the Brain Scaffold Builder (BSB) (https://bsb.readthedocs.io/en/latest/), a neuroinformatic framework allowing a detailed microcircuit reconstruction based on neuron morphologies and orientations and the incorporation of active neuronal and synaptic properties (De Schepper et al., 2022). The SNN was made of ~3×10^4^ extended-Generalised Leaky Integrate and Fire (E-GLIF) neurons (Geminiani et al., 2018, 2019a) and ~1.5×10^6^ alpha-shaped conductance-based synapses (Roth and van Rossum, 2013). The SNN simulations were performed using NEST (Plesser et al., 2015; Jordan et al., 2019)

#### 2.1.1 Neuron model

The E-GLIF formalism describes the time evolution of membrane potential (V_m_) depending on two intrinsic currents to generate slow adaptation (I_adap_) and fast depolarisation (I_dep_), using a the system of three ODEs (Geminiani et al., 2018)

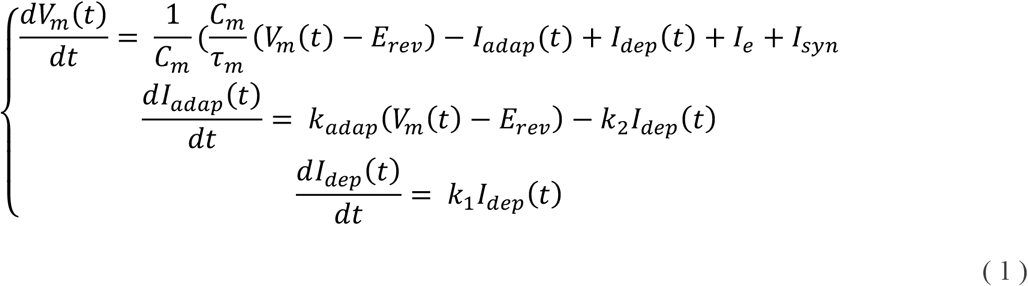

where I_syn_ = synaptic current (it models the synaptic stimulus, see section 2.1.2); C_m_ = membrane capacitance; τ_m_ = membrane time constant; E_rev_ = reversal potential; I_e_ = endogenous current; k_adap_ and k_2_ = adaptation constants; k_1_ = decay rate of I_dep_. When a spike occurs, state variables are updated as follows:

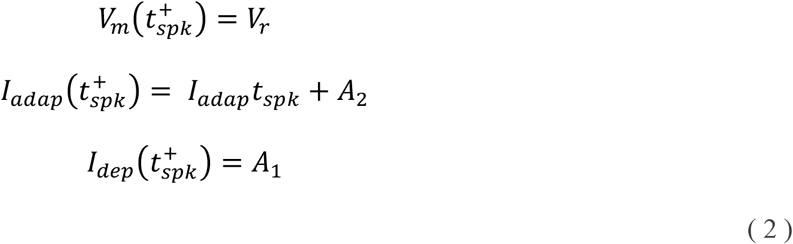

where 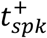 = time instant immediately following the spike time t_spk_; V_r_ = reset potential; A_2_, A_1_ = model currents update constants. E-GLIF models were implemented using parameter sets specific for each neuronal population (Geminiani et al., 2019a) as shown in Table 1.

**Table 1.** Neuron parameters.

| Parameter | Name | unit | GrC | GoC | MLI | PC |
| --- | --- | --- | --- | --- | --- | --- |
| $g_L$ | Leak conductance | nS | 0.29 | 3.30 | 1.60 | 7.10 |
| $C_m$ | Membrane capacitance | pF | 7.00 | 145.00 | 14.60 | 334.00 |
| $\tau_{ref}$ | Refractory time | ns | 1.50 | 2.00 | 1.59 | 0.50 |
| $\tau_m$ | Membrane time constant | ns | 24.15 | 44.00 | 9.12 | 47.00 |
| $E_L$ | Resting potential | mV | -62.00 | -62.00 | -68.00 | -59.00 |
| $V_{th}$ | Threshold potential | mV | -41.00 | -55.00 | -53.00 | -43.00 |
| $V_r$ | Reset potential | mV | -70.00 | -75.00 | -78.00 | -69.00 |
| $k_{adap}$ | Adaptation constant | MH<sup>-1</sup> | 0.02 | 0.22 | 2.03 | 1.50 |
| $k_2$ | Adaptation constant | ms<sup>-1</sup> | 0.04 | 0.02 | 1.10 | 0.04 |
| $k_1$ | Decay rate | ms<sup>-1</sup> | 0.31 | 0.03 | 1.89 | 0.19 |
| $A_2$ | Update constant | pA | -0.94 | 170.01 | 5.86 | 172.62 |
| $A_1$ | Update constant | pA | 0.01 | 259.99 | 5.95 | 157.62 |
| $I_e$ | Endogenous current | pA | -0.89 | 16.21 | 4.45 | 891.04 |

#### 2.1.2 Synaptic model

Connections between neural populations were modelled as conductance-based synapses:

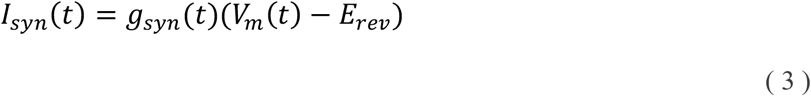

When a spike occurs, the conductance g_syn_ changes according to an alpha function:

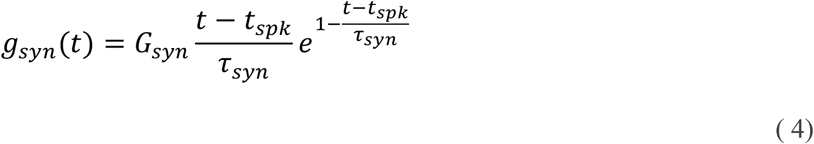

where G_syn_ is the maximum conductance change and τ_syn_ the synaptic time constant. E-GLIF neuron models and conductance-based synaptic models used in SNN simulations provided the functional reference of cerebellar spiking activity for MF development.

### 2.2 MF design

The design of the cerebellar multi-layer MF was based on the extensive knowledge of cerebellar anatomy and physiology summarized in previous cerebellar cortex network models (Geminiani et al., 2018, 2019c; De Schepper et al., 2021).

#### 2.2.1 Architecture

The cerebellar MF included the main neuronal populations of the cerebellar cortex - GrC, GoC, MLI and PC (Figure 2) and the corresponding excitatory and inhibitory synapses. The MF network topology reproduced the multi-layer organisation of the cerebellar cortex. Granular layer at the cerebellar input stage includes GrC and GoC receiving external input (*ν*_drive_) from mossy fibers. GrC excite GoC, which, in turn, inhibit themselves and GrC forming recurrent loops. GrC represent the excitatory input for the molecular layer constituted by MLI and PC. MLI inhibit PC, which are the sole output of the cerebellar cortex and shape the deep cerebellar nuclei activity through inhibition.

**Figure 2.**
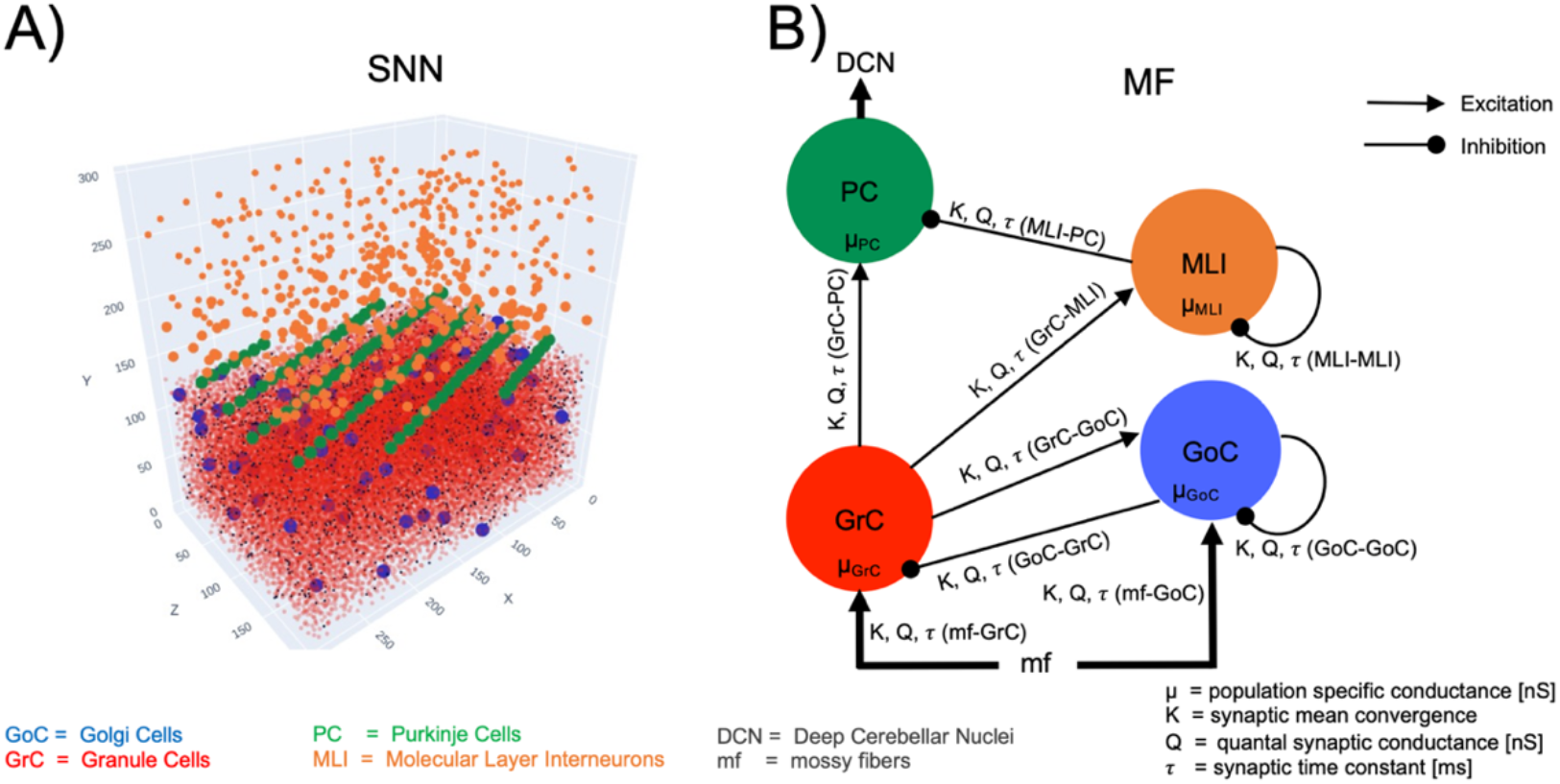
Multi-layer MF architecture and parameters. **A**: Spiking Neural Network model (SNN). The cerebellar cortical volume (length = 300, width = 200, height = 295 μm3) contained a total of 29230 neurons including 28615 GrC, 70 GoC, 446 MLI and 99 PC. **B**: Multi-layer MF architecture with neuronal populations connected according to anatomical knowledge. The main cerebellar neuron types are included: GrC and GoC, receiving input from mossy fibers, MLI and PC, which are the sole output of the cerebellar cortex. Each population receive excitatory and/or inhibitory input activity *ν* from presynaptic populations, depending on their specific conductance μ and on the synaptic properties of each connection (convergence, synaptic conductance, and time constant).

Although other neurons have been reported to play a role in the cerebellar microcircuit (e.g. Lugaro cells (Melik-Musyan and Fanardzhyan, 2004), and unipolar brush cells (Mugnaini et al., 2011) for the sake of simplicity we have limited the present model to the canonical architecture that is thought to generate core network computations.

To connect the nodes of the MF network, synaptic parameters were set according to those of the reference SNN (Table 2). The connection probability for each connection type (K) was derived from the convergence ratio in a cerebellar cortical volume (De Schepper et al., 2022). The quantal synaptic conductance and synaptic time decay (Q, *τ*) was derived from the weights and time constants of the corresponding synapse models (Geminiani et al., 2019a).

**Table 2.** Presynaptic parameters.

| Presynaptic connection | K | Q [nS] | $\tau$ [ms] | $E_{\text{rev}}$ [V] |
| --- | --- | --- | --- | --- |
| mf-GrC | 4.00 | 0.230 | 1.9 | 0 |
| GoC-GrC | 3.50 | 0.240 | 4.5 | -80 |
| mf-GoC | 57.10 | 0.240 | 5.0 | 0 |
| GrC-GoC | 501.98 | 0.437 | 1.25 | 0 |
| GoC-GoC | 2592.00 | 0.007 | 5.0 | -80 |
| GrC-MLI | 243.96 | 0.154 | 0.64 | 0 |
| MLI-MLI | 1418.69 | 0.005 | 2.0 | -80 |
| GrC-PC | 489.16 | 0.510 | 1.1 | 0 |
| MLI-PC | 10.28 | 1.244 | 2.8 | -80 |

Parameters used to set up the inter-population connectivity of the multi-layer MF cerebellar network. The parameters were extracted from the spiking neural network simulating the cerebellar cortex spiking activity (Geminiani et al., 2019b). Mf = mossy fibers, GrC = Granule Cells, GoC = Golgi Cells, MLI = Molecular Layer Interneurons (Basket cells and Stellate cells). K = pre-synaptic connectivity resulting by weighting the mean synaptic convergence with the number of synapses; Q = quantal synaptic conductance in nS; $\tau$ = synaptic decay time constant; $E_{\text{rev}}$ = reversal potential that is 0 V for excitatory synaptic connections and -80 V for inhibitory synaptic connections.

#### 2.2.2 TF Computation

The TF is defined for each population as a mathematical construct that takes the activity of the presynaptic population (*ν*_s_) as input and provides an average population activity signal as output (*ν*_out_) (Kuhn et al., 2004).

A purely analytic derivation of the TF (using approximations and stochastic calculus, e.g. Brunel & Amit, 1997) was not possible given the relative complexity of the neuronal (E-GLIF) and synaptic (alpha-waveform) models considered here. Therefore, the TFs presented here relied on a semi-analytical approach that couples an approximate analytical estimate with an optimization step to capture the firing response of analytically intractable models (Zerlaut et al., 2016, see also Brunel & Sergi, 1998 for a similar approach). More details can be found in Zerlaut et al. (2016) but we summarize the approach below.

The analytical template for the TF (indicated with F in the equations for sake of simplicity) of all neuron types is derived from the probability to be above threshold in the fluctuation-driven regime (Kuhn et al., 2004):

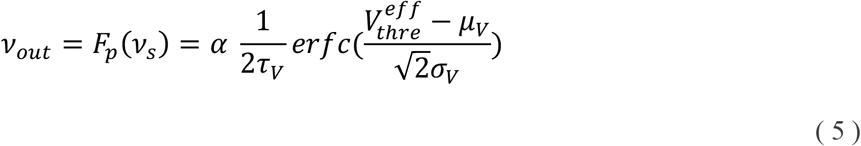

where *erfc* is the error function while μ_V_, σ_V_^2^ and τ_V_ are the average, variance and autocorrelation time respectively of the membrane potential fluctuations. Two phenomenological terms were introduced: 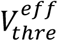, an effective firing threshold to capture the impact of single cell non-linearities on firing response (Zerlaut et al., 2016) and α, a multiplicative factor to adapt the equations also to high input frequency regimes (Carlu et al., 2020). Those two terms were optimized for each neuron type (steps *d* and *e* respectively) from single neuron simulations of input-output transformation in terms of firing rate (i.e., the numerical TF, see step b). The TF depends on the statistical properties of the subthreshold membrane voltage dynamics (mean = μ_V_, standard deviation = σ_V_^2^ and autocorrelation time τ_V_, calculated in step *c*). These in turns depend on the average population conductances that are computed with the biologically-grounded functional parameters derived from SNN models at single neuron resolution (step *a*), bringing the physiological properties into the MF mathematical construct.**Errore. Il segnalibro non è definito**.

##### a) Equations of Population-specific conductance

For each neuronal population, the average conductance was defined as a function of the presynaptic inputs, according to the topology described in **section 2.1** (Figure 3.2.):

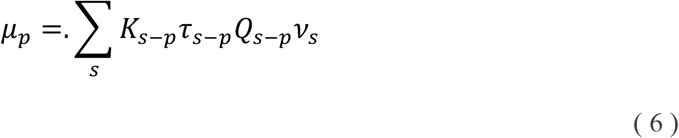

where, for each population *p* (*p* = GrC, GoC, MLI, PC), K_s-p_, *τ*_s-p_, Q_s-p_ are the connection probabilities, synaptic decay times and quantal conductances of the connection for each presynaptic population *s* (e.g., for *p* = GrC, *s-p* is mossy fiber-GrC and GoC-GrC), *v*_s_ is the presynaptic population activity in Hz computed as explained in (b).

##### b) Numerical TF

The reference functional target was the neuronal spiking activity obtained in SNN simulations (*in vivo* conditions) as described in **section 2.1**. The activity of GrC, GoC, MLI and PC embedded in the SNN was simulated for different input amplitudes (*ν_drive_*) in the range 0-80 Hz, with input spikes generated from a Poisson distribution. For each *ν_drive_*, the simulation lasted 5 seconds with a time resolution of 0.1 ms. The working frequencies of each population were extracted by averaging the spiking neuron firing rates. For each population, the outcome of numerical TF computation was a template of dimension equal to the number of presynaptic populations, resulting in 2D numerical TFs for GrC, MLI and PC and 3D numerical TF for GoC (Figure 3A).

**Figure 3.**
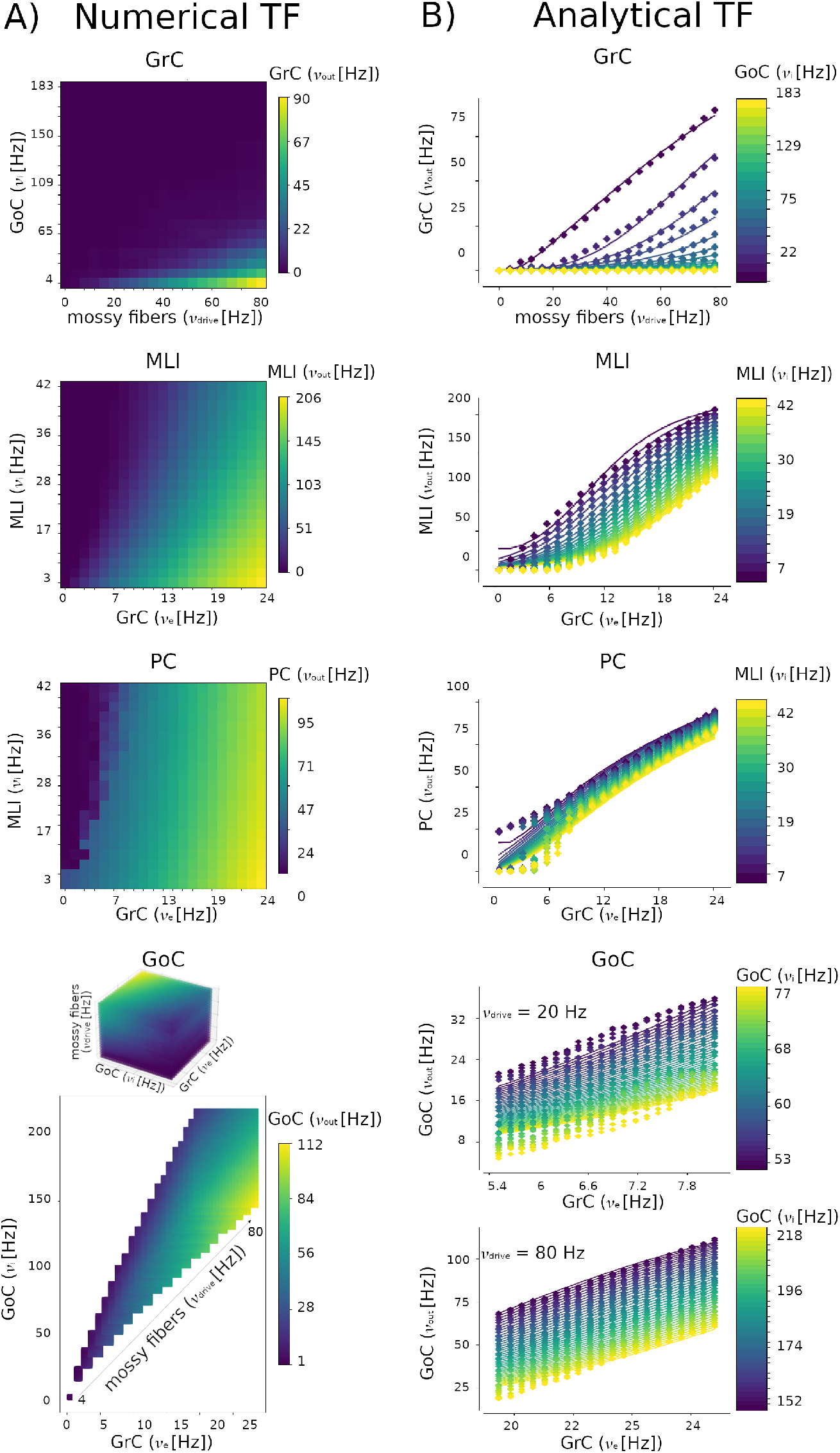
Numerical TFs and the corresponding Analytical fitting. The simulation used to compute the numerical TFs last 5 seconds, with a time step = 0.1 ms. **A)** A 2D numerical TF template is reported for GrC, MLI and PC, which receive inputs from two presynaptic populations. A 3D numerical TF template is reported for GoC, which receive input from 3 presynaptic sources, i.e., mossy fibers, GrC and GoC. From the 3D domain of frequencies combination, only the physiological working frequencies of GrC and GoC are considered for mossy fibers inputs from 0 Hz to 80 Hz, as obtained in corresponding spiking neural network simulations and 3D numerical TF for GoC which receive three presynaptic inputs. **B)** The numerical TFs are used to fit the corresponding analytical TF. For each neuronal population, the average activity obtained in spiking neural network simulations is represented in color code, for each combination of input activity levels. The 2D analytical TF presents a non-linear trend, while the GoC Analytical TF shape is almost linear both for lower *ν*_drive_ = 20 Hz and higher *ν*_drive_ = 80 Hz.

The 2D numerical TF of each population was computed as the population firing rate when receiving the firing rates of the presynaptic populations, given a certain *ν_drive_*: for example, for the numerical TF numerical template of PC, the average firing rates of MLI and GrC were computed for each *ν_drive_* in the range 0-80 Hz. Then, these quantities were used as presynaptic signals to stimulate PC and a numerical template was obtained from the resulting PC firing rate, for each combination of presynaptic activities). The 3D TF numerical template of GoC was computed following the same strategy but considering 3 presynaptic signals (GrC, GoC, mossy fibers). GrC excitation and GoC self-inhibition were extracted from SNN simulations and the mossy fibers excitation corresponded to *ν_drive_*.

##### c) Statistical moments of the MF

The statistical moments included in the MF are μ_V_, σ_V_ and τ_V_. Starting from the conductances of the presynaptic populations, the average conductance *μ_G_* of the target population reads:

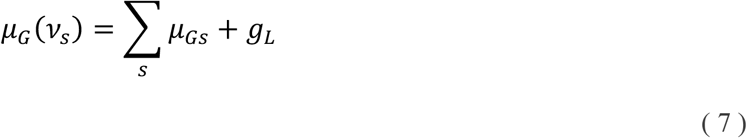

Where *μ_Gs_* is the presynaptic population conductance (equation (4)) and *g_L_* is the leak conductance of the target population (Table 1). Then, the effective membrane time constant of the target population is computed from *μ_G_* as:

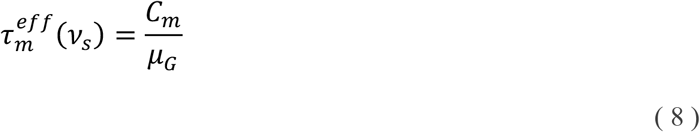

Where *C_m_* is the membrane capacitance (Table 1). The first statistical moment, i.e., the average of membrane potential fluctuation *μ_V_* reads:

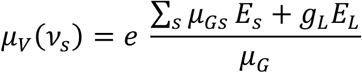

With E_s_ = reversal potential of the presynaptic connection (0 mV for the presynaptic excitatory populations and −80 mV for the presynaptic inhibitory populations), E_L_ = rest potential of the target population (Table 1).

This expression is adapted from Zerlaut et al. 2018 to model the alpha synapses consistently with the models used in the SNN (equations (1), (2), and (3)). Consequently, the variance and the autocorrelation time of membrane fluctuations result in:

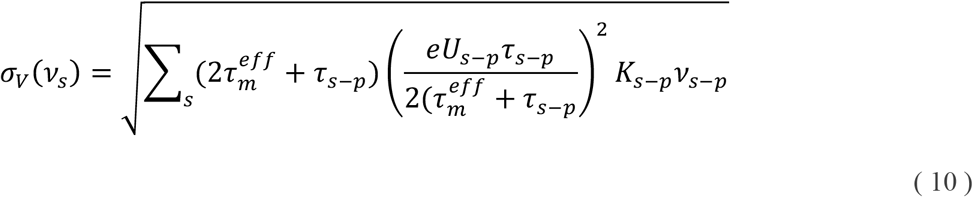

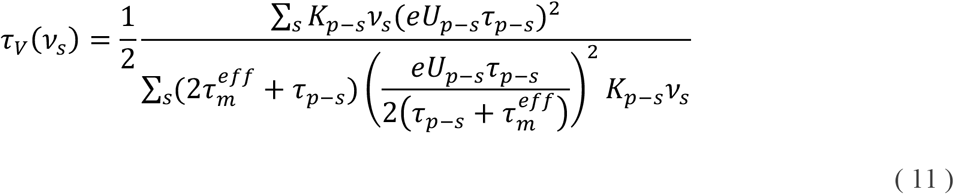

With 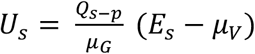

##### d) Phenomenological threshold

The ability of the analytical template (5) to capture different firing behavior is given by the introduction of 5 parameters in the phenomelogical threshold term. The phenomenological threshold is expressed as a linear combination of the *V_m_* fluctuations properties whose coefficients are linearly fitted to the numerical TF data (Zerlaut et al., 2016):

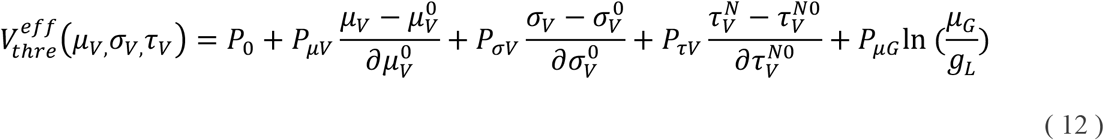

Where 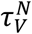 is *τ_V_* adjusted with the ratio between membrane capacitance and leak conductance 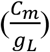, and 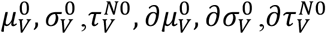 are rescaling constants to normalize the contribution of each term (Zerlaut et al., 2018). P are the polynomial coefficients which are the target of the fitting procedure to compute the analytical TF as explained in e) (see Table 3).

**Table 3.** Fitted coefficients of the Analytical TFs.

**Fitted coefficient P [V]**
|  | **P<sub>0</sub>** | **P<sub>μV</sub>** | **P<sub>σV</sub>** | **P<sub>τV</sub>** | **P<sub>μG</sub>** |
| --- | --- | --- | --- | --- | --- |
| **GrC** | -0.426 | 0.007 | 0.023 | 0.482 | 0.216 |
| **GoC** | -0.144 | 0.003 | 0.011 | 0.031 | 0.011 |
| **MLI** | -0.128 | -0.001 | 0.012 | -0.093 | -0.063 |
| **PC** | -0.080 | 0.009 | 0.004 | 0.006 | 0.014 |

P coefficients computed with the fitting procedure explained in Zerlaut et al., 2016 and extended to E-GLIF neurons.

##### e) Analytical TF

The statistical moments in equations (9), (10), (11) and the phenomenological threshold in equation (12) were plugged into equation (13) and the phenomenological threshold is computed through a fitting procedure described in Zerlaut et al. 2016. The TFs specific for the cerebellar populations are reported in Figure 3B.

The parameter alpha (equation (5)) was set to an optimal value for each population to fit both low and high frequencies (Carlu et al., 2020). The analytical TF, together with the statistical moments *μ_V_*, *σ_V_*, and *τ_V_* defined the cerebellar MF equations.

#### 2.2.3 Multi-layer equations

The multi-layer MF was developed as a set of equations capturing the interdependence of the population-specific TFs, tailoring the isocortical MF described in (El Boustani and Destexhe, 2009) for excitatory-inhibitory networks to the cerebellar network. This formalism describes the network activity at a time resolution T which is set to ensure a Markovian dynamic of the network: T should be large enough to ensure memoryless activity (e.g., it cannot be much lower than the refractory period, which would introduce memory effects) and small enough so that each neuron fires statistically less than once per time-bin T. The choice of T is quite crucial and here it was tailored to account for cerebellar dynamics as explained in.

The model describes the dynamics of the first and the second moments of the population activity for each population. The cerebellar network was build up with four interconnected populations (GrC, GoC, MLI, PC) receiving external input from mossy fibers (mf) (Figure 2), thus resulting in twenty differential equations: the four population activities (*ν*_GrC_(t), *ν*_GoC_(t), *ν*_MLI_(t), *ν*_PC_(t)) and the driving input (*ν*_mf_ (t) = *ν_drive_*(t)), the four variances of the population activities (*c*_GrC-GrC_(t), *c*_GoC-GoC_(t), *c*_MLI-MLI_(t), *c*_PC-PC_(t)) and the one of the driving input from mossy fibers (c_mf-mf_(t)), the six covariances among population activities (*c*_GrC-GoC_(t), *c*_GrC-PC_(t), *c*_GrC-MLI_(t), *c*_GoC-MLI_(t), *c*_GoC-PC_(t), c_MLI-PC_(t)) and the four covariances between population activities and the driving input (*c*_GrC-mf_(t), *c*_GoC-mf_(t), *c*_MLI-mf_(t), *c*_PC-mf_(t)). Einstein’s notation was used to report the differential system in a concise form:

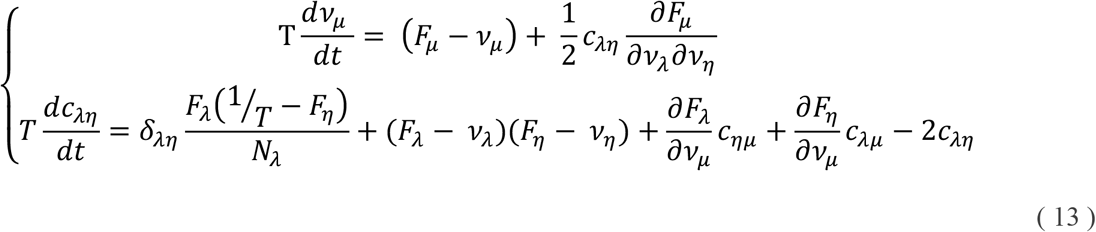

Where *ν*_μ_ is the activity of population μ; c_λ*η*_ is the (co)variance between population λ and *η*; N is the number of cell included in population λ. According to Einstein’s notation, a repeated index in a product implies a summation over the whole range of values. TF dependencies on the firing rate of presynaptic populations are omitted yielding F_μ_ instead of F_μ_(*ν*_s_) with μ = {GrC, GoC, MLI, PC} and *s* is the presynaptic population (e.g. *F_GoC_* = *F_GoC_* (*ν_drive_*, *ν_GrC_*, *ν_GoC_*)). The model equations (13) were numerically solved using forward Euler method with an integration step of 0.1 ms.

#### 2.2.4 Timing Optimisation

The MF time-constant T was optimised by comparing the model prediction with experimental data of the cerebellar Granular layer (Figure 4). The simulated average activity was interpolated with the experimental Local Field Potential (LFP) measured with high-density microelectrode arrays (HD-MEA) in the granular layer of acute mouse cerebellar slices (Mapelli and D’Angelo, 2007)

**Figure 4.**
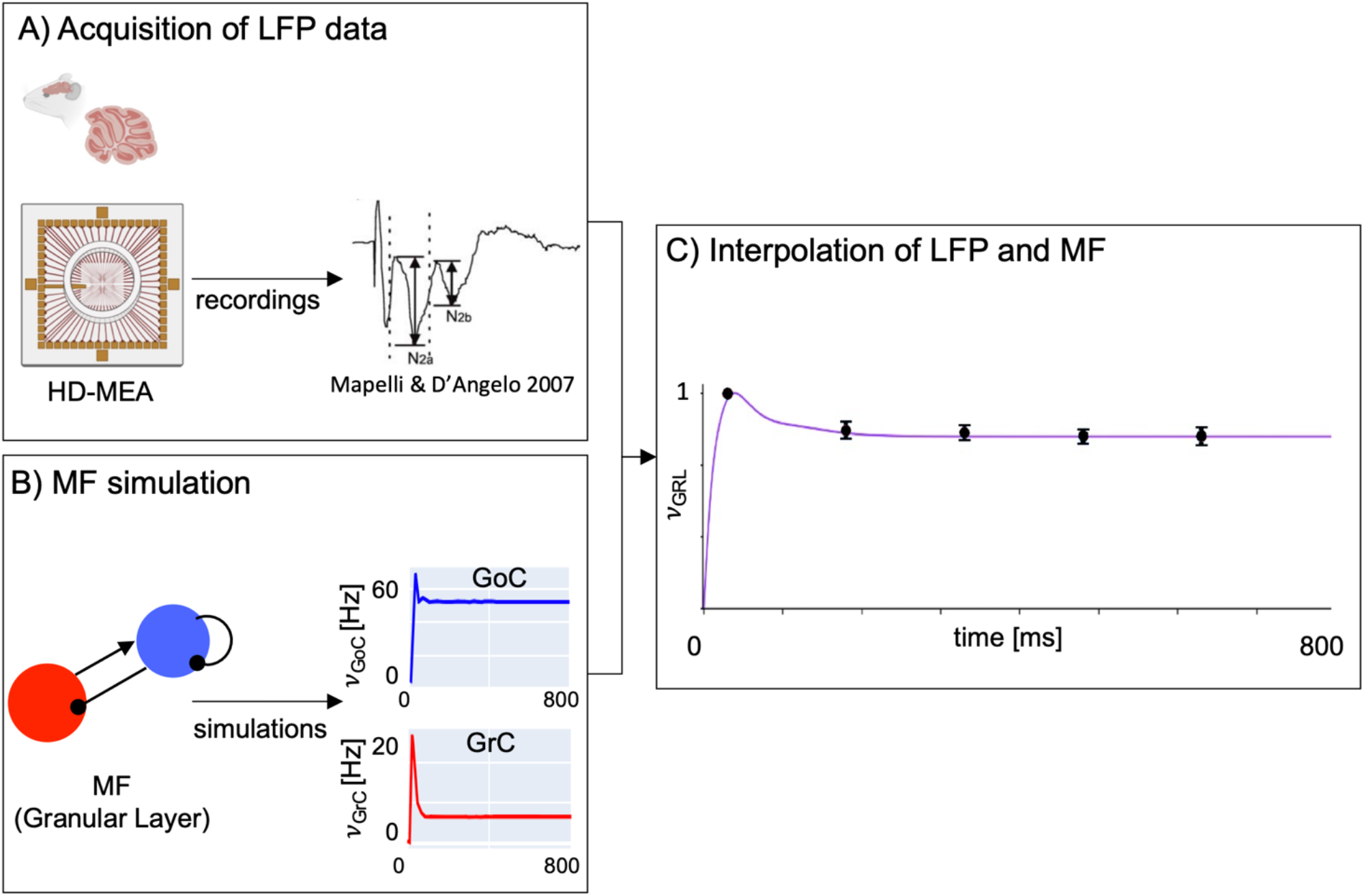
The mean-field time constant. **A)** Experimental acquisition of LFP. LFP signals were acquired in the cerebellar granular layer of acute mice cerebellar para-sagittal slices using HD-MEA in response to five stimulation pulse trains of 50 Hz (Unpublished data, courtesy of Lisa Mapelli and Anita Monteverdi). **B)** MF simulation. The activity of the granular layer was simulated with the cerebellar MF using a stimulation protocol emulating experimental LFP recordings. **C)** Interpolation of LFP and MF. The weighted average of the predicted Granular Layer activity (*ν*_GRL_, violet line) interpolates the LFP data (mean ± SD; dots and bars). The relative weights of GoC and GrC are 13% and 87%, respectively. The optimal T value is 3.5 ms ± 5%, (mean ± mean absolute error between *ν*_GRL_ and LFP).

LFP data were recorded at 37 °C. The external stimulus consisted in a pulse train of 5 stimuli of 50 Hz amplitude. This stimulation protocol was repeated nine times changing the HD-MEA recording channels across each experiment (Figure 4A). The LFP signals recorded were averaged across the nine experiments resulting in five values that represented the average of each pulse of the input trains. These average records were normalised on the amplitude of the signal recorded after the first stimulus.

The cerebellar MF simulation protocol was configured with a *ν_drive_* = 50 Hz for 100 ms, reproducing the experimental protocol (Figure 4B). A range of plausible T values was evaluated according to literature (El Boustani and Destexhe, 2009; Zerlaut et al., 2016, 2018; Carlu et al., 2020). MF simulations were performed with a systematic change of T value and the granular layer average activity was calculated by a weighted-mean of GrC and GoC activity. The weight of GrC and of GoC was computed as the ratio of the spiking surfaces (GoC/GrC) resulting in 0.13 (Mapelli and D’Angelo, 2007). The granular layer average activity was normalised on the maximum peak, and it was interpolated with LFP recordings (Figure 4C). For each simulation the mean absolute error was computed to select the T value that minimised the discrepancy with the LFP signals. Since the granular layer is the driving layer of the network, the optimal T value was extended to the molecular and Purkinje layers.

### 2.3 Constructive and Functional validity

For constructive and *functional validity*, the cerebellar MF was tested using stimulation protocols designed to assess its ability in reproducing proper cerebellar dynamics and stimulus-response patterns.

Four different stimulation protocols were defined, each lasting 500 ms:

i. *ν_drive_* = Step function. A square wave with steps of amplitude 50 Hz, and lasting 250 ms to reproduce a conditioned stimulus (e.g. a sound)
ii. *ν_drive_* = Theta-band sinusoid. A sinusoidal input with rate amplitude set at 20 Hz and frequency at 6Hz (theta band) to simulate the whisker movements experimental conditions.
iii. *ν_drive_* = Combination of alpha, theta, and gamma band sinusoid. A combination of 3 sinusoidal inputs, with fixed rate amplitude at 40 Hz and frequency at 1Hz, 15Hz and 30Hz respectively reproducing a EEG-like pattern.
iv. *ν_drive_* = Step function plus multi-band sinusoid. Summation of the step function described in (i) with amplitude of 20 Hz and sinusoidal function including alpha, theta and gamma band with the same frequency of (iii) and amplitude of 7.5 Hz to simulate a more complex input like a conditioned stimulus overlapped to a realistic basal activity

Population activities predicted by the MF were overlayed to the Peristimulus Time Histogram (PSTH) with bin = 15 ms, computed from spiking activities of the SNN in NEST simulations of the same stimulation protocols. Then, the PC activity, i.e., the output of the cerebellar cortex, was quantified as mean ± standard deviation of the firing rate for both MF and SNN simulation. Boxplots were shown to quantitively compare MF and SNN outcomes, and the computational efficiency of each model was measured as computational time in seconds required for each simulation performed with MF and with NEST.

### 2.4 Predictive Validity

For predictive validity, MF parameters were tuned to explore the MF sensitivity to modifications of local mechanisms. These modifications were derived from experimental studies on neural correlates of behavior in functional or dysfunctional conditions, focusing on inhibitory control and long term plasticity on PCs (Wulff et al., 2009; ten Brinke et al., 2015)

#### 2.4.1 MLI feed forward inhibition modulation

Feedforward inhibition from molecular layer interneurons regulates adaptation of PCs. Impact of MLI-PC conductance on PC activity was explored by defining different values of MLI-PC synaptic strength W_MLI-PC_ = [5, 30, 100, 150, 200, 250] %, where W_MLI-PC_ = 100% represents the standard condition, rates lower than 100% model disinhibited activity, while rates higher than 100% extra-inhibition.

W_MLI-PC_ was added to the Analytical TF_PC_ as a modulatory parameter of the presynaptic input *ν*_MLI_, resulting in a modulation of MLI contribution in PC population conductance (35) defined as follows

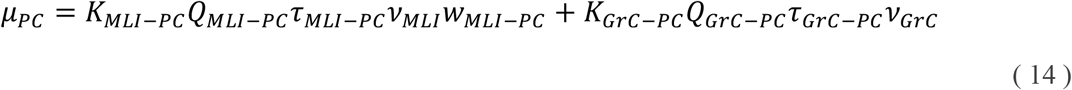

Each simulation lasted 500 ms with a 50Hz driving input of 250 ms after 125 ms of resting.

The Area Under Curve (AUC) of PC activity, PC peak and the depth of the pause were computed as quantitative scores for each W_MLI-PC_ value AUCs and PC peaks were normalised on the respective values corresponding to the standard condition defined as W_MLI-PC_ = 100%.

#### 2.4.2 PC Learning

Long term potentiation and depression (LTP and LTD) are forms of synaptic plasticity at the basis of brain learning processes (Bliss and Cooke, 2011). In the cerebellum, motor learning is driven by PC activity modulation, regulated by the plasticity of the synapses between parallel fibers (pfs – from GrC) and PC, resulting in a reduction of PC activity due to LTD and in an increase of PC activity due to LTP (Mittmann and Husser, 2007; Prestori et al., 2013). To simulate pf-PC plasticity, the following synaptic strengths (w_pf-PC_) were explored to investigate the consequent PC modulation: [5, 20, 35, 50, 65, 80, 100, 120, 135, 150, 165, 180, 200, 235, 250, 265] % where 65% corresponds to the decrease of pf-PC strength during motor learning according to animal experiments value (ten Brinke et al., 2015), and the others were defined to capture the trend of pf-PC plasticity mechanism. The values minor than 100% means LTD occurred, while the values higher than 100% represent LTP occurrence. The strategy applied was analogous to equation (13).

PC AUC and PC peak were computed for each w_pf-PC_ value as quantitative score to correlate the amount of LTD and LTP with the output activity of the cerebellar cortex.

### 2.5 Hardware and software

The SNN was built with the BSB release 3.0 (https://bsb.readthedocs.io/en/v3-last) and the numerical simulations were performed with NEST version 2.18 (https://zenodo.org/record/2605422).

The MF design, the timing optimisation, the MF validation, and the MF predictive simulations were implemented in Python 3.8. Functions packages written for the present work are available on *https://github.com/RobertaMLo/CRBL_MF_Model*.

All optimisation procedures and simulations were run on a Desktop PC provided with AMD Ryzen 7 2700X CPU @ 2.16GHz with 32 GB RAM in Ubuntu 16.04.7 LTS (OS).

## 3 Results

### 3.1 The cerebellar MF

The workflow for reconstructing the cerebellar MF is shown in Figure 1 leading to a condensed representation using 4 neuronal populations for GrC, GoC, PC, MLI neurons (Figure 2). The MF was designed based on structural and functional parameters extracted from SNN simulations and the time constant was optimized LFP experimental data. The MF working frequencies were extracted from NEST simulations of the cerebellar SNN exploring multiple *ν_drive_* from 4 to 80 Hz. Then these frequency ranges were used to set different plausible presynaptic signals in defining the Numerical TFs of each population (Figure 3.3A). The ranges were [0.42, 24.17] Hz for GrCs, [3.63, 183.15] Hz for GoCs, and [3.27, 41.66] Hz for MLIs. Note that PC working frequencies were not computed since PC activity is only projected forward to the cerebral cortex, therefore PCs never play the role of presynaptic population in this cortical cerebellar microcircuit. The α parameters that maximised the fitting performance for each population were α _GrC_ = 2, α _GoC_ = 1.3, α _MLI_ = 5 and α _PC_ = 5. The fitted coefficients P are reported in Table 3

2D Analytical TFs show a sigmoidal trend in relation with excitatory inputs (Figure 3B). GoC inhibition strongly affects the GrC Analytical TF; for *ν*_GoC_ higher than 100 Hz, GrC Analytical TF is almost 0 Hz. For low inhibition, e.g., *ν*_GoC_= 13 Hz, GrC Analytical TF is almost linear. MLI Analytical TF presents a well-defined sigmoidal trend depending on *ν*_GrC_ and modulated by the auto-inhibition, with resulting activity frequency spanning from 0 Hz up to 200 Hz. PC Analytical TF presents an increasing trend ranging from 0 to 100 Hz in relation to *ν*_GrC_ from 0 to 25 Hz, with the modulation due to the inhibitory control from MLIs. 3D Analytical TF of GoC shows a linear trend both for low and high *ν_drive_*.

The cerebellar MF resulted in a set of 20 second order differential equations including the specific population TFs, where 4 equations described the time variation of population activity, and the remaining 16 equations modelled the covariances of the interconnected populations. Figure 4 shows the result of T optimisation: for T = 3.5 ms the average granular layer activity (purple line) interpolates the experimental LFPs (red dots) with a mean absolute error of 3%.

### 3.2 Constructive and functional validity

The validation of the cerebellar MF was obtained generating neuronal population dynamics with different stimulation protocols and comparing them with the corresponding SNN simulations (Figure. 3.5). For all the protocols, the simulation lasted 500 ms and was performed with the hardware and software specified in **section 2.5**.

#### Step function (i)

The MF was tested with a 250 ms@50 Hz step function on the mossy fibers, simulating a conditioned stimulus (Figure 3.5A) (Jirenhed and Hesslow, 2011). GrC activity rapidly raised at the beginning of the step input, then strongly decreased due to inhibitory GoC activity. GoCs, after an initial small peak due to both the direct incoming input and the GrC excitation, maintained a steady-state activity for all the step duration. The dynamics of GrC-GoC interplay faithfully reproduced the feedback loop between GoCs and GrCs. GrC was the excitatory input for the molecular layer, and both MLI and PC activity arose in correspondence of the GrC initial peak. Thus, exploiting network di-synaptic delays, MLIs reduced PC activity soon after its maximum, generating the typical burst-pause pattern of these neurons. The PC pause is due to both the single neuron parameters and to inhibitory local connectivity in the microcircuit. After this rapid transient, the activity of MLIs and PCs reached a steady-state. In the MF, fast dynamics at the input step onset and at the steady-state matched SNN simulations for all neuronal populations.

#### Theta band sinusoid (ii)

Simulated dynamics of all cerebellar populations showed oscillations driven by the input reproducing whisker movements (Figure 3.5B) (Popa et al., 2013). GrC activity projected to the molecular layer a sinusoidal-shaped signal at 0.05-5Hz, contributing to an oscillatory behaviour in GoCs, and causing an oscillation in MLI between at 23-41 Hz, and in PC activity at 42-69 Hz. Oscillations had comparable amplitude in MF and SNN simulations and occurred in the same frequency ranges (except for MLI activity that was slightly higher in MF that SNN).

#### Alpha, gamma and theta band sinusoid (iii)

Combination of alpha, gamma and theta sinusoids (with EEG-like frequency (Del Percio et al., 2017) of 1 Hz, 15 Hz, 30 Hz, respectively) caused an irregular oscillation in the input carried by mossy fibers (2-47 Hz range) (Figure 3.5C) Oscillations had comparable amplitude in MF and SNN simulations and occurred in the same frequency ranges.

#### Step function plus multi-band sinusoid (iv)

The summation of repeated step function (i) and multi-band sinusoid (iii) resulted in an irregular input (Figure 3.5D) depicting in-vivo noisy baseline activity with a conditioned stimulus superimposed. GrC activity faithfully transmitted the driving input, with peaks at ~21 Hz. In correspondence with the GrC excitatory peak, MLIs peaked at ~130 Hz and PCs at ~100 Hz.

The responses had comparable dynamics and amplitude in MF and SNN simulations.

### 3.3 Predictive validity

The PC response to a 50Hz step-stimulus was described by a peak at 97 Hz followed by a pause down to 68 Hz; then a steady-state of 78 Hz was attained. MLI-PC and pf-PC modulation (Figure 3.6) perturbed this reference condition.

#### 3.3.1 MLI-PC feed forward inhibition

Inhibitory interneurons control the generation of burst-pause patterns in PC, which is fundamental for shaping the cerebellar output during motor learning (Casali et al., 2019; Kim and Augustine, 2021). For instance, knock-out of MLI inhibition on PC impacts on vestibulo-ocular reflex adaptation (Wulff et al., 2009). Here in MF simulations, when the MLI-PC conductance was reduced to 5% of the reference condition, the burst-pause dynamics of PC was lost, so that the PC firing settled directly back to baseline (which was elevated due to lack of inhibitory control). Conversely, when PCs were over-inhibited by the MLIs (MLI-PC conductance increased to 250% of reference condition), the pause was deeper. The PC overall activity (AUC), and PC Peak reveal an exponential trend that decays for higher MLI-PC conductances. The PC pause shows a decreasing sigmoidal trend for higher MLI-PC conductances (Figure 3.6A).

#### 3.3.2 PC plasticity

Long Term Depression (LTD) and Long Term Potentiation (LTP) at pf-PC synapses are though to drive cerebellar adaptation and learning. The overall activity and the peak of PCs showed a linear and a sigmoidal trend, respectively, with the increase of pf-PC weight. With decreased pf-PC strength, the peak was reduced or disappeared, and the steady-state activity reached lower levels. With increased pf-PC strength, both the peak and steady-state values were increased (Figure 3.6B). During a typical cerebellum-driven behaviour, the eyeblink classical conditioning (EBCC), a level of suppression of about 15% has been reported and correlated with a stable generation of associative blink responses at the end of the learning process (ten Brinke et al., 2015). SNN simulation got the same result by setting AMPA-mediated pf-PC synapses = 35% (De Schepper et al., 2022). In MF simulations, for w_pf-PC_ = 65%, which corresponded to a reduction of pf-PC conductance of 35%, PC activity presented a 22% reduction of the peak and a 10% reduction of the AUC, falling into the experimental range of PC suppression (ten Brinke et al., 2015).

## 4. Discussion

This work shows, for the first time, a MF of cerebellar cortex. According to its bottom-up nature, the MF transfers the microscopic properties of neurons (including GrC, GoC, MLI, and PC) and synapses of the cerebellar cortex into a condensed representation of neural activity through its two main statistical moments, mean and variance. The construction and validations strategies adopted here make the present MF an effective representation of the main physiological properties of a canonical cerebellar module (D‘Angelo and Casali, 2013)

### 4.1 MF design and validation

#### The cerebellar network and TF formalism

The cerebellar cortex MF was based on the same general formalism previously developed for the isocortex MFs (Moran et al., 2013; Di Volo et al., 2019; Carlu et al., 2020; Huang and Lin, 2021). However, the cerebellar cortex MF benefitted of a previously validated SNN to precisely remap cellular and synaptic biophysical properties and network topology. Moreover, the electrophysiological properties of neurons were represented using non-linear EGLIF models and the synapses with alpha-based conductance functions. This resulted in three main advantages.

First, the parameters of population specific TFs were validated against biophysically detailed models of neurons and the connectome was derived from precise scaffold model reconstructions providing a direct link to the biological microcircuit (De Schepper et al., 2022).

Secondly, the equations of μ_V_, *σ*_V_ and *τ*_V_ included in the TF formalism were adapted to model the alpha-shaped synapses and to maintain rise-times in synaptic dynamics. For comparison, previous MFs (Zerlaut et al., 2016, 2018) used exponential synapses, which provide a less realistic approximation due to their instantaneous rise time (Brette and Gerstner, 2005).

Finally, MF included 4 different species of neurons that were modelled using either 2D or 3D TFs to account for the multiplicity of their inputs (Figure 3). It is worth noting that the analysis of both 2D TFs of GrC, MLI and PC, and 3D TF of GoC were fitted considering only physiological input combinations computed from single-neuron computational models. In the fitting procedures, indeed, fine-tuned parameters were included to maintain a strong physiological correspondence. The 3D dimension of the GoC TF avoided to merge the excitatory input from GrC and mossy fibers (*ν*_GrC_ and *ν_drive_*, respectively), enabling us to investigate distinct excitatory input contributions to granular layer dynamics and to the whole cerebellar MF. By fixing the excitatory mossy fibers driving input, we assessed the power of the Analytical TF in simulating spiking network activity for inputs at both low and high frequency (e.g., see Figure 3.4 with *ν_drive_*= 20 Hz and = *ν_drive_*= 80 Hz).

A technical issue incurred while fitting the numerical TF. The TF formalism models the difference between phenomenological threshold 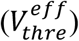 and population average responses (μ_V_) through the complementary error function (erfc in equation (5)), providing an immediate interpretation of how single neuron activity was related with statistics of population dynamics (μ_V_, *σ*_V_, *τ*_V_). Since erfc is stiff and limited between −1 and 1, it may not accurately follow the numerical TF distribution at the boundaries, losing precision at high frequencies. This problem was circumvented by tuning the parameter alpha (Carlu et al., 2020), making the TF analytical expression reliable over the whole range of input working frequencies.

#### MF tuning

The inclusion of precise structural and functional parameters in the design of cerebellar MF (see Figure 2 and 3) generated a biology grounded model that could be validated at a higher scale with the prediction of cerebellar dynamics (multi-layer equation 13). The dynamics of the cerebellar cortex are several times faster compared to those of the cerebral cortex (D’Angelo, 2011), so that the MF time constant, T, must be optimized accordingly. The MF time constant was optimized using experimental LFP recordings from the cerebellar granular layer acquired on the same spatial scale of the MF. The best fitting was obtained by accounting for the smaller contribution of GoC than GrC activity (13% vs. 87% (Dieudonné, 1998; D’Angelo et al., 1999)) to LFPs (Mapelli and D’Angelo, 2007), revealing that the MF time constant of the cerebellar cortex is T=3.5 ms with mean absolute error of 3% (Figure 3.4). T is definitely smaller than in cerebral cortex MFs, which range up to 20 ms (Zerlaut et al., 2018; Di Volo et al., 2019; Carlu et al., 2020), and captures the peculiar high speed of cerebellar dynamics (D’Angelo, 2011). This result further confirms the need of a MF specifically tailored on the cerebellum functional and topological parameters.

The optimal T value was plugged into equation 13 resulting in a second order differential equation system of interdependent TFs capturing the dynamics of multiple cerebellar populations and their covariances. This rich pool of equations allowed our cerebellar MF to reproduce a variety of cerebellar dynamics in response to different inputs (see **section 2.3** and shown in Figure 5) which, by comparison with the equivalent SNN output, provided the benchmark for constructive and functional validity. A rapidly changing input like a step function reproduced a conditioned stimulus (Jirenhed and Hesslow, 2011) carried by the peripherical mossy fibers, causing rich dynamics in the cerebellar cortex including the typical PC burst-and-pause responses (Herzfeld et al., 2015). Adding a multi-sinusoidal input to the conditioned stimulus replicated a more physiological condition accounting for background activity. This resulted in rich PC dynamics, which still maintained burst-and-pause responses. A sinusoidal input was meant to emulate more complex experimental conditions, like those determined by whisker movement (Popa et al., 2013; Yamazaki and Igarashi, 2013; Antonietti et al., 2017; Masoli et al., 2020a; Gagliano et al., 2021, 2022). In particular, a multi-sinusoidal waveform (comprised of frequencies in the EEG spectrum) was used to emulate composite inputs from the cerebral cortex (Del Percio et al., 2017; Tzvi et al., 2022).

**Figure 5.**
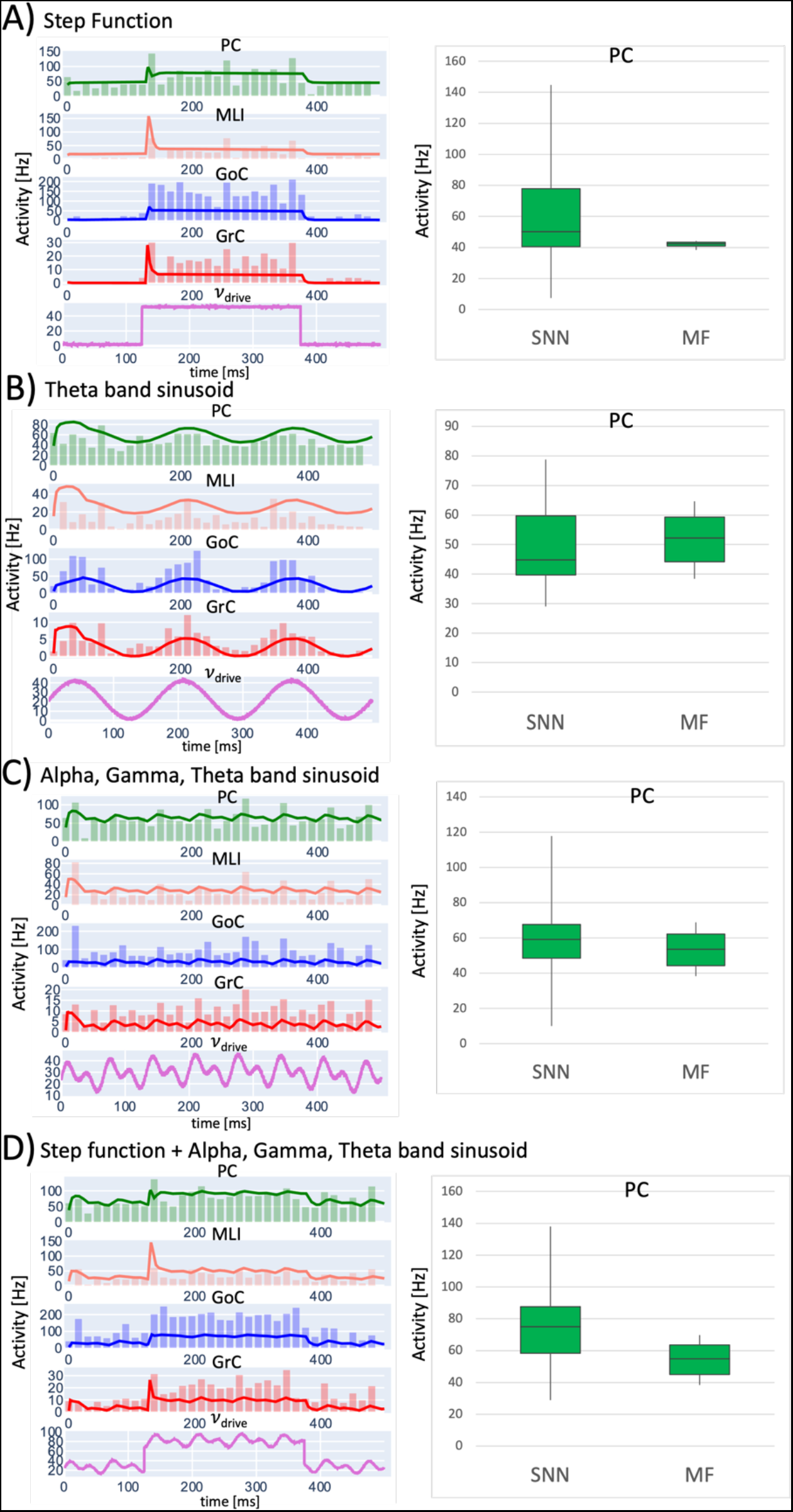
Constructive and functional validity. Comparison of SNN and MF activity in cerebellar cortical populations in response to different driving input (*ν_drive_*) patterns, lasting 500 ms with time resolution = 0.1 ms. **A)** step function (Conditioned stimulus-like); **B)** sinusoidal input in the theta band (whisker-like); **C)** multi-frequency sinusoidal function (EEG-like); **D)** combined stimulus summing (A) step function and (C) multi-frequency sinusoidal. The trace of MF activity is overlayed to the spiking activity, which is represented as a PSTH (time bins of 15ms). In all cases, MF activity is within physiological ranges, capturing also fast changes of activity due to instantaneous input changes in step-function input protocols. The boxplots of PC simulated activity with SNN and MF, shows that the MF is able to respond to the different stimulation patterns within the same frequency ranges of SNN.

The cerebellar MF reproduced the known aspects of circuit physiology revealed experimentally. Indeed, GrCs respond to impulsive inputs with bursts curtailed by GoCs and are also able to follow slower input fluctuations (D’Angelo and De Zeeuw, 2009). The PCs generate burst-pause responses that are accentuated by MLIs (Masoli and D’Angelo, 2017; Rizza et al., 2021). The cerebellar MF quantitatively reproduced these patterns matching the corresponding SNN simulations, with the only exception of the maximal GoC responses, which did not increase as expected with rapidly changing inputs like the step function. This is a consequence of the lack of GoC heterogeneity in MF, in which all GoC are collapsed in a homogeneous population despite their biological heterogeneity (Galliano et al., 2010). Except for this, MF predicted with good approximation the SNN cerebellar output, i.e., the PC activity (Boxplot in Figure 5). The inclusion of probabilistic kernels addressing parameter heterogeneity could also help improving the MF fitness (Di Volo and Destexhe, 2021).

### 4.2 MF predictions

A critical step in model validation is to demonstrate its ability to predict functional states not used for model construction. The cerebellum is well known for the ability to change its network functioning because of synaptic and non-synaptic plasticity. However, different from SNN, the MF did not include plasticity mechanisms. Thus, we directly tested the MF ability to predict the effects of plasticity expression by mapping a set of precomputed synaptic changes on the MF itself. MF reproduced the impact of MLI-PC conductance confirming that, also in the cerebellar MF, the complex burst-pause behaviour of PC is tuned through the MLI-PC connectivity.

The cerebellar MF reproduced the experimental recordings in EBCC experiments on behaving mice (ten Brinke et al., 2015) pointed out a PC LTD of 10% in terms of overall activity and 22% for the peak (Figure 6 w_pf-PC_ = 65%). This protocol corresponds to a reduction of 35% of AMPA-mediated pf-PC in SNN simulation, therefore our MF capability of capturing synaptic mechanism is further validated against in-vivo recordings.

**Figure 6.**
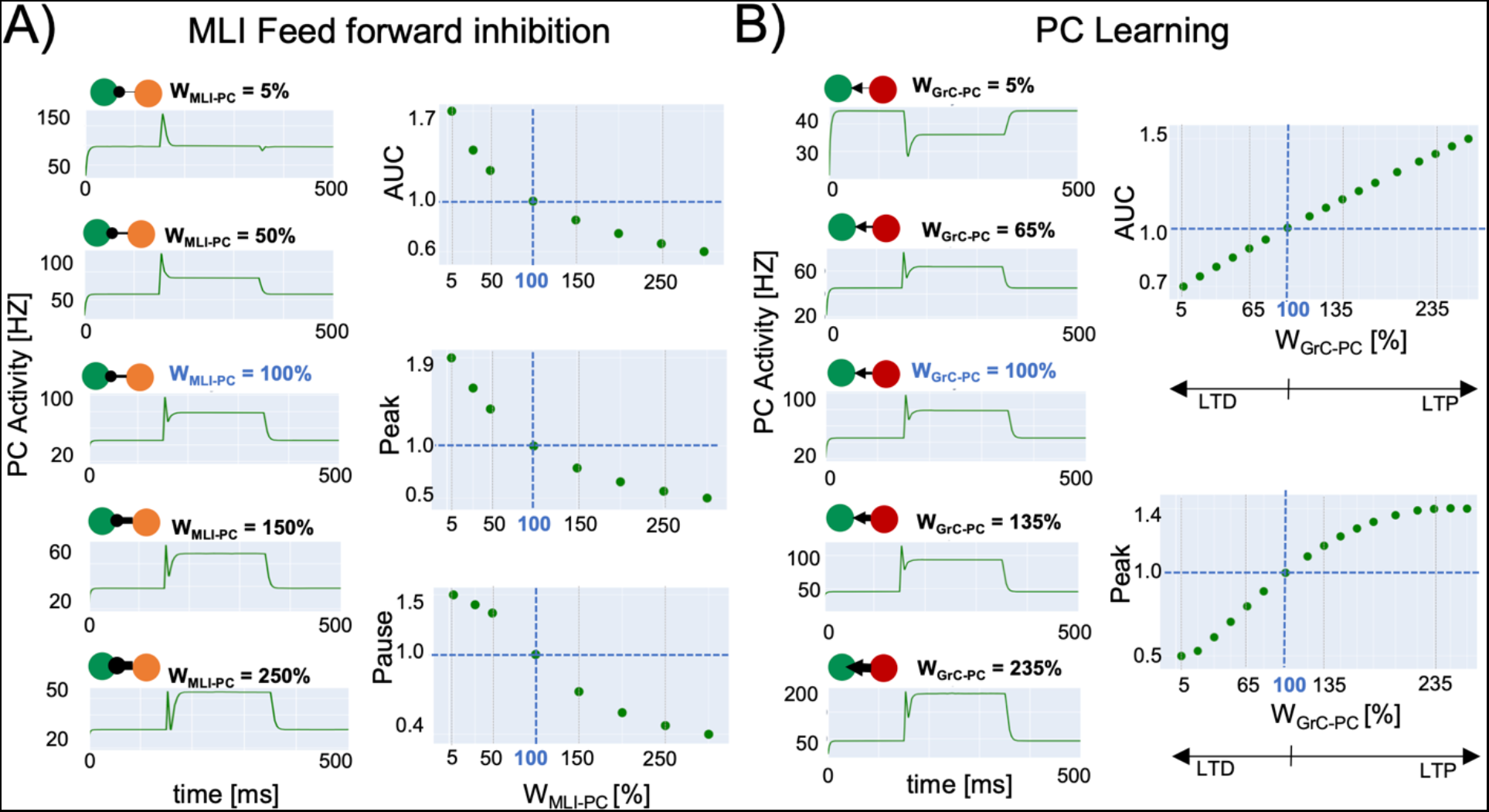
Predictive validity. MF simulations using different strengths at PC connections. Simulations last 500 ms with a time resolution of 0.1 ms and *νdrive* is a step at 50 Hz lasting 250 ms starting at 125 ms. Quantitative score normalized to baseline (w = 100% in blue). **A) MLI-PC feed forward inhibition**. The change of total activity (measured as AUC) and the initial peak amplitude decrease exponentially with the MLI-PC strength. The amplitude of the pause after the peak response decreases with MLI-PC strength following an inverse sigmoidal function. **B) PC Learning** with different GrC-PC plasticity conditions (LTP and LTD). AUC linearly increases with the GrC-PC strength, matching the experimental values in experimental learning protocols. The initial peak caused by the step stimulus onset increases with GrC-PC until saturation, following a sigmoidal function.

In aggregate, the cerebellar MF, despite a lower level of detail than SNN, was able to reproduce complex physiological mechanisms and predict the activity changes caused by synaptic modulation (ten Brinke et al., 2015; De Schepper et al., 2022). Therefore, this MF is a flexible tool that can be used to investigate advanced physiological or pathological properties by tuning the input of TF on a target population (TF_PC_ in these simulations), without complicating neither the TF fitting procedure nor the model equations. The procedure of parameters tuning might pave the way for further manipulation to remap physiological and/or pathological features onto the MF. Identifying and extracting biophysical meaningful features from subject-specific data, like diffusion weighted imaging or spectroscopy, might allow the MF tailoring onto subject-specific characteristics and to combine functional simulations with (micro)structural information.

### 4.3 Performance vs. realism

The MF approximated a complex SNN of ~3×10^4^ neurons and ~1×10^6^ synapses (**section 2.1**) with 20 equations reducing the computational time by 60% with lower memory requirements. Nevertheless, TFs fitting could be improved replacing the procedure in **section 2.2.2** with a lookup table-based algorithm, which might yield to a gain in computational time up to an order of magnitude. This will represent a definite advantage when performing long-lasting simulations reflecting the acquisition time of *in vivo* recordings like EEG and fMRI or when simulating learning processes in closed-loop controllers. On the other hand, thanks to its bottom-up nature it was possible to maintain the biological realism in responses to various stimuli, including simulations of learning-induced firing rate changes and pathological conditions at the neuronal population level, obtaining a good balance between computational load and biological plausibility. This will allow to make predictions on the underlying neural bases of ensemble brain signals and to identify the elementary causes of signal alterations in pathology.

### 4.4 Conclusions and perspectives

In aggregate, the cerebellar cortex MF enforces a bottom-up approach tailored to the specific structural and functional interactions of the local neuronal populations and has a substantial constructive and functional validity. By accounting for a variety of representative patterns of discharge in cerebellar cortical neurons, the present MF can be considered as an effective proxy of the biological network. The internal model parameters inform about the average properties and variance of fundamental mechanisms in the circuit, namely intrinsic and synaptic excitation, and can therefore be used to remap biological properties onto the MF (Naskar et al., 2021). In future applications, this will allow to tune the MF toward specific functional or dysfunctional states that affect the cerebellum. Among these it is worth mentioning ataxias (Pedroso et al., 2019; Rosenthal, 2022), paroxysmal dyskinesia (Mendonça and Alves da Silva, 2021; Ekmen et al., 2022), dystonia (Mahajan et al., 2021; Morigaki et al., 2021), autistic spectrum disorders (Bruchhage et al., 2018; Kelly et al., 2020) as well as other pathologies like and multiple sclerosis (Tornes et al., 2014; Schreck et al., 2018), dementia (Monteverdi et al., 2022) and Parkinson disease (Wu and Hallett, 2013; Shen et al., 2020), in which a cerebellum involvement has been reported. The cerebellar MF could be applied to whole-brain simulators using TVB and DCM, as much as it has been done before for the isocortical MF in TVB (Pinotsis et al., 2012; Goldman et al., 2019; Sadeghi et al., 2020; Ruffini and Deco, 2021). Considering the specificity of signal processing in different brain regions, this approach represents a definite step ahead compared to the classical one adopting generic neural masses for all brain regions. This is a promising active field in theoretical neuroscience and clinics, indeed applications of specific MF to Parkinson disease have been already implemented by reconstructing a model specifically tailored on basal-ganglia and connected with thalamocortical circuit (BGTCS MF), accurately reproducing parkinsonian state (van Albada and Robinson, 2009; van Wijk et al., 2018). Integrating BGTCS and cerebellar MF, and in general MF specifics of brain regions embedded into the motor circuit, would remarkably improve the simulation of brain dynamics, allowing to compare dysfunctional oscillations with physiological activity.

On the theoretical side, TVB simulations using classical neural masses (Wong and Wang, 2006) for all brain nodes (Palesi et al., 2020; Monteverdi et al., 2022) can now be compared to those using the cerebellum MF. At the other extreme of the spectrum, TVB with embedded cerebellar MF can be compared to TVB-NEST co-simulations (Meier et al., 2022), in which spiking neurons are represented explicitly (Geminiani et al., 2018, 2019b, 2019a; De Schepper et al., 2022). These comparisons will inform us about the impact of populations, neurons, and spikes on ensemble brain dynamics and whole brain computations (D’Angelo and Jirsa, 2022).

In conclusion, the cerebellar MF represents the first step toward a new generation of models capable of bearing biological properties into virtual brains that will allow to simulate the healthy and pathological brain towards the overarching aim of a personalized brain representation and ultimately personalized medicine and the technology of brain digital twins (Amunts et al., 2022).

## Author Contribution

FP, CC, CGWK, and ED’A conceptualized the study. CC and ED’A coordinates the study. RL and AG designed the pipeline and RL performed the analyses. RL, AG, ED’A wrote the paper. YZ and AD provided support and guidance with data analysis and interpretation. All authors reviewed and contributed to the article and approved the submitted version.

## Acknowledgment

We thank Robin De Schepper for useful discussion on Brain Scaffold Builder framework, Alessio Marta for an insight into the mathematical formalism of mean field, and Lisa Mapelli and Anita Monteverdi for sharing unpublished local field potential recordings.

## Funding

This research has received funding from the European Union’s Horizon 2020 Framework Program for Research and Innovation under the Specific Grant Agreement No. 945539 (Human Brain Project SGA3) to ED, CGWK, FP and AD, and under the Marie Sklodowska-Curie grant agreement No. 892175 to YZ. CGWK received funding from BRC (#BRC704/CAP/CGW), MRC (#MR/S026088/1), Ataxia UK, MS Society (#77), Wings for Life (#169111). CGWK is a shareholder in Queen Square Analytics Ltd.This research has also received funding from Centro Fermi project “Local Neuronal Microcircuits” to ED. Special acknowledgement to EBRAINS and FENIX for informatic support and infrastructure.

## Notes

### Competing Interest Statement

The authors have declared no competing interest.

